# Functional anatomy inspires recognition of Ediacaran *Dickinsonia* and similar fossils as Mollusca (Coleoidea) and precursors of Cambrian Nectocaridids and extant cuttlefish and squid

**DOI:** 10.1101/057372

**Authors:** Bernard L. Cohen

## Abstract

A functional interpretation of the problematic Ediacaran fossils *Podolimurus* and similar organisms such as *Dickinsonia* indicates that they are hitherto unrecognized members of Mollusca: Coleoidea and precursors of Cambrian Nectocaridids and of extant cuttlefish and squid. This interpretation enables a new understanding of their taphonomy and reveals how this has deceived previous attempts to understand them.

## Introduction

The difficulty of correctly identifying the affinities of fossil organisms results from a combination of factors including the mists of time, the generally uncertain and incomplete conditions and processes that lead to fossilisation and, for the most ancient forms, the unfamiliarity of their morphology. For Ediacaran soft-bodied organisms the difficulties are particularly extreme, resulting in many taxa being described as *Problematica*. Such mysteries led me to study the recent description of *Podolimurus mirus* (Dzik & Martyshyn, 2015, see especially fig. 4b) and to realise that a functional interpretation of the described anatomy is available and that it yields an improved interpretation of this and similar organisms (e.g. *Dickinsonia*) as precursors of extant squid and cuttlefish.

## Methods and Materials

The analysis is literature-based, cited papers (and others referred to but not cited) being downloaded from journal web-sites and printed at high resolution. Further details are not necessary.

## Results and Discussion

*Podolimurus* and similar *Problematica* have been interpreted as possessing a bilaterally symmetrical, serially repeated, dorsal quilt for which no clear functional interpretation has been provided. Unusually, some specimens of this taxon also reveal an anterior structure, interpreted as intestinal diverticula (Dzik & Martyshyn, 2015). This structure has not been reported in *Dickinsonia*, because it is absent, has decayed away, or is hidden.

Brief consideration of the possible functional roles of the dorsal quilt suggests that it is a buoyancy aid, and this idea immediately suggests that it is the soft-tissue homologue (and precursor) of the bilaterally symmetrical, serially repeated chambers of the cuttle-bone found in extant coleoids and the analogue (or homologue) of the gas-filled flotation chambers of nautiloids and ammonoids. The physiology of buoyancy regulation in the extant coleoid *Sepia* has been well studied (Denton, Gilpin-Brown & Howarth, 1961; Denton & Gilpin-Brown, 1961). Whereas the osmotically active membrane of the cuttlebone surrounds the entire structure, the many quilt compartments of these *Problematica* may have independently controlled the osmotic composition of their internal medium, greater surface area potentially enabling faster equilibration to depth changes that those measured in *Sepia* (Denton et al., 1961; Sherrard, 2000). So far as we are aware no other, and no more plausible, functional role has been proposed for the quilted structure in any fossil, and certainly none that has a well-studied extant analogue or homologue.

Reinterpreting the quilt as a buoyancy aid enables the taphonomy of the parent organisms to be differently and better understood. Consistent with the impressions they made in the fossil-bearing strata, the walls of the quilt’s cells were relatively rigid structures, presumably pressure-resistant and impermeable in life (perhaps cartilaginous), and therefore less liable to microbial decay than other, softer tissues. Therefore (if present) head, eyes, tentacles, mantle siphon (jet), gut, ventral mantle wall, gills and gonads are all likely to have been removed by predation, autolysis and/or microbial action in the water column such that, except in special circumstances, only the quilt is likely to reach the bottom and become a candidate for burial (e.g. Kear, Briggs & Donovan, 1995). Thus, preservation will have been highly selective and biased. It may therefore be predicted that well-preserved organisms of these taxa will rarely be found. Intact animals are likely to reach the sea-bottom only in shallow waters, but these are likely to be well-oxygenated, and therefore most favourable to decay. Well-preserved fossils are likely to be formed only exceptionally, when live animals have been caught in anoxic conditions that also inhibit anaerobic decay, and this is followed by rapid burial (e.g. under Burgess Shale-type conditions).

Many proposals have been made for the biology and phyletic relationships of *Dickinsonia*, the most prominent of which envisages it feeding like a placozoan on benthic microbial mats (Sperling & Vinther, 2010). This proposal suffers from serious defects. For example, that placozoans are still highly enigmatic with, until very recently very few being known and little being known for certain about their life history or feeding mode. Thus inferences of Ediacaran biology based upon extant placozoans are inevitably weak and stretched, especially by the required assumption that *Dickinsonia* has some ill-defined feeding mechanism on its ventral side. More significantly, extant placozoans have no specific, morphological synapomorphies that directly indicate affinity with *Dickinsonia*, *Poldolimurus* or any other taxon. Given the newly identified functional role of the quilt, the supposed “feeding traces” of *Dickinsonia* can now be understood as nothing more than the result of largely decomposed corpses settling repeatedly on the shore before being stranded permanently by an outgoing tide and subsequently buried under sand carried by an incoming tide. Furthermore, in the deposit from which *Podolimurus* fossils were recovered microbial mats are uncommon (Dzik & Martyshyn, 2015) suggesting that there is no justification for assuming a specific association between them and the fossil-forming taxon.

### Unexpected, independent confirmation of the proposed affinity

Following recognition of the probable coleoid affinity of *Dickinsonia*-like organisms a search was made for further relevant literature, and this revealed the existence of Nectocaridids, already recognised as stem-group cephalopod-like organisms (Smith, 2013; Smith & Caron, 2010). Despite some differences, age, preservation style, taphonomy and morphology nevertheless allow *Nectocaris* to be interpreted with some confidence as a member of the same clade as *Dickinsonia* and *Podolimurus*. It is possible that the so-called “bars” of Nectocaridids, for which no clear function has been proposed, are the walls of their bilateral buoyancy quilts, but the match is not clear and the taphonomy of *Nectocaris* (Smith, 2013, fig. 1) suggests that this taxon may instead have regulated its buoyancy by dynamic or chemical lift, as do extant non-Sepioid coleoids (Packard, 1972). Active buoyancy control is an obvious adaptive asset in a nectonic predator, allowing the neutral flotation depth to be varied to permit energy-efficient swimming at different depths and in different salinities.

### Implications for metazoan evolution

1. Molluscan diversification clearly occurred in the Ediacaran (see e.g. Stöger, Kano, Knebelsberger, Marshall, Schwabe & Schrödl, 2013). It is therefore very likely that brachiopod and annelid stem groups diverged from their molluscan sister-group and other lophotrochozoan stem taxa much earlier than generally envisaged and certainly long before the start of the Cambrian.

2. Coleoids may be the longest-lasting and most successful of all nectonic predators, having early “invented” buoyancy-control mechanisms that allows energy-efficient swimming at different depths.

## Acknowledgements

I am grateful to Alekandra Bitner and Carsten Lüter for commenting on a draft and to Dave Sherratt for advice.

